# Optimization of a *Cannabis sativa* micropropagation protocol (chemotype III) to preserve its cannabinoid profile

**DOI:** 10.1101/2025.05.26.656187

**Authors:** María Micaela González, Gabriel Yañuk, Analía Sannazzaro, Estefanía Butassi, Melina Di Liberto, Eliane Perez, Mónica Hourcade, Laura Svetaz, Marina Clemente

## Abstract

A micropropagation protocol was developed using axillary buds of the Charlie’s Dream variety of *Cannabis sativa* (chemotype III, THC≪CBD), under the hypothesis that clonal propagation preserves the CBD/Δ9-THC profile between donor and micropropagated plants. The effects of plant growth regulators BAP and TDZ (0.5, 1, 2.5 μM) in MS medium with or without vitamin supplementation (MS vs. MSS) were determined. The progress in the growth of the axillary buds was evaluated by analyzing the shoot length, the number of shoots per explant, and the foliar area of the shoots. The role of naphthaleneacetic acid (NAA) and indolebutyric acid (IBA) in the rooting process was also evaluated *in vitro* and *ex vitro*. Resins were extracted from dried flowers of donor and micropropagated plants using cold ethanol treatment. The CBD and Δ9-THC contents were analyzed by GC/MS. The results show that vitamin supplementation in the MS medium did not improve any of the growth parameters evaluated. However, significant differences were observed when comparing TDZ and BAP. At 15 days post-initiation of micropropagation (dpim) from axillary buds, the explants treated with 0.5 and 1 μM TDZ showed a greater number of shoots per explant and larger foliar area, compared to those treated with any concentration of BAP. However, by 30 dpim, explants treated with BAP showed an improved performance, reaching shoot numbers and foliar area values similar to those observed with TDZ. It is worth mentioning that no differences were observed in the shoot length at all concentrations analyzed for both hormones. Likewise, the explants grown in 2.5 μM TDZ showed greater callus production and vitrification. The effects of supplementation with the hormone gibberellin (7 µM) or exposure of the shoots to red light were also evaluated. The results indicated that combining these treatments with BAP at 0.5 and 1 µM significantly enhanced shoot elongation compared to combinations with TDZ at the same concentrations. On the other hand, *in vitro* rooting was unsuccessful. However, direct rooting in soil was observed, and root length was further enhanced when shoots were pretreated with 2.5 μM IBA. The CBD/Δ9-THC ratio remained stable between donor and micropropagated plants, supporting the development of standardized micropropagation protocols. Our results confirm that clonal micropropagation is a reliable method to preserve cannabinoid profiles in *C. sativa*.

## 1. Introduction

*Cannabis sativa* is mainly dioecious and allogamous, resulting in high genotypic and phenotypic variability [1]. This inherent variability hinders the consistent production of uniform, high-quality plants from seed that comply with regulatory and market standards [2]. For this reason, cannabis for medicinal use has been propagated through clonal methods. Traditionally, clonal propagation is performed by cutting female plants with high cannabinoid levels. Generally, large quantities of plants can be produced from a single donor plant using this propagation technique, as the cannabis plant is relatively easy to root [3]. However, this procedure involves allocating between 10 and 15% of the ground space for a single commercial operation. Furthermore, under these cultivation conditions, donor plants are susceptible to attack by insects, viruses, fungi, and bacteria that can be transmitted to the cuttings and seriously affect the production and traceability of the crop. This is of great relevance in cannabis, as there are currently very few pathogen control options registered for cultivation, and, in addition, there is a strong preference for pesticide-free consumption [4,5].

*In vitro* micropropagation represents an alternative approach to conventional clonal propagation, enabling large-scale multiplication under strictly controlled, aseptic conditions. The sterile environment inherent to this technique facilitates the production of pathogen-free plantlets, thereby reducing biotic stress [6]. In the clonal micropropagation of *C. sativa*, a primary challenge is to develop robust methods that ensure reproducibility of protocols across different genotypes and chemotypes [7]. To date, most micropropagation protocols for the cannabis plant are based on the multiplication of shoots from apical and axillary nodes. The most widely adopted and effective approaches involve the use of commercial plant growth regulators (PGRs) [6,8–10]. Nonetheless, variability in morphogenic responses among cultivars remains a challenge hindering efficient and reliable production [6]. Likewise, due to genetic and varietal diversity within *C. sativa*, bottlenecks persist during the regeneration, growth, and rooting stages of *in vitro cultivation*, limiting the development of rapid, reliable, and efficient production systems. Therefore, it is crucial to investigate and optimize *in vitro* culture protocols to establish standardized micropropagation methods tailored to each chemotype and/or genotypic/phenotypic variety. In this work, we aimed to optimize the *in vitro* culture conditions of the *C. sativa* variety Charlie’s Dream (chemotype III, THC<<CBD) using axillary buds, based on the hypothesis that clonal micropropagation preserves chemotype characteristics while maintaining consistent CBD and Δ9-THC levels between the donor and their micropropagated counterparts.

## 2. Materials and Methods

### 2.1. Plant material

Seeds of the Charliés Dream variety (chemotype III, THC<<CBD) of *C. sativa* were purchased commercially and germinated on filter paper moistened with distilled water inside Petri dishes in a culture chamber at 23°C±2°C in the dark. Germinated seeds were sown in pots containing a substrate composed of a mixture of perlite and soil (1:1) and grown in a culture room maintained at 23°C±2°C, under a 22-h light/2-h dark photo-period and a photon flux density of 200 µmol m^-2^ s^-1^ with LED lighting.

### 2.2. Explants

The evaluated explants (∼1 cm in length) comprised axillary and apical buds from two donor plants and four micropropagated plants originating from one of these donors (Fig. 1a). The outer tissue was cut with a scalpel and two sets of primordial leaves were kept to protect the meristem from excessive damage by disinfection.

**Figure 1.**
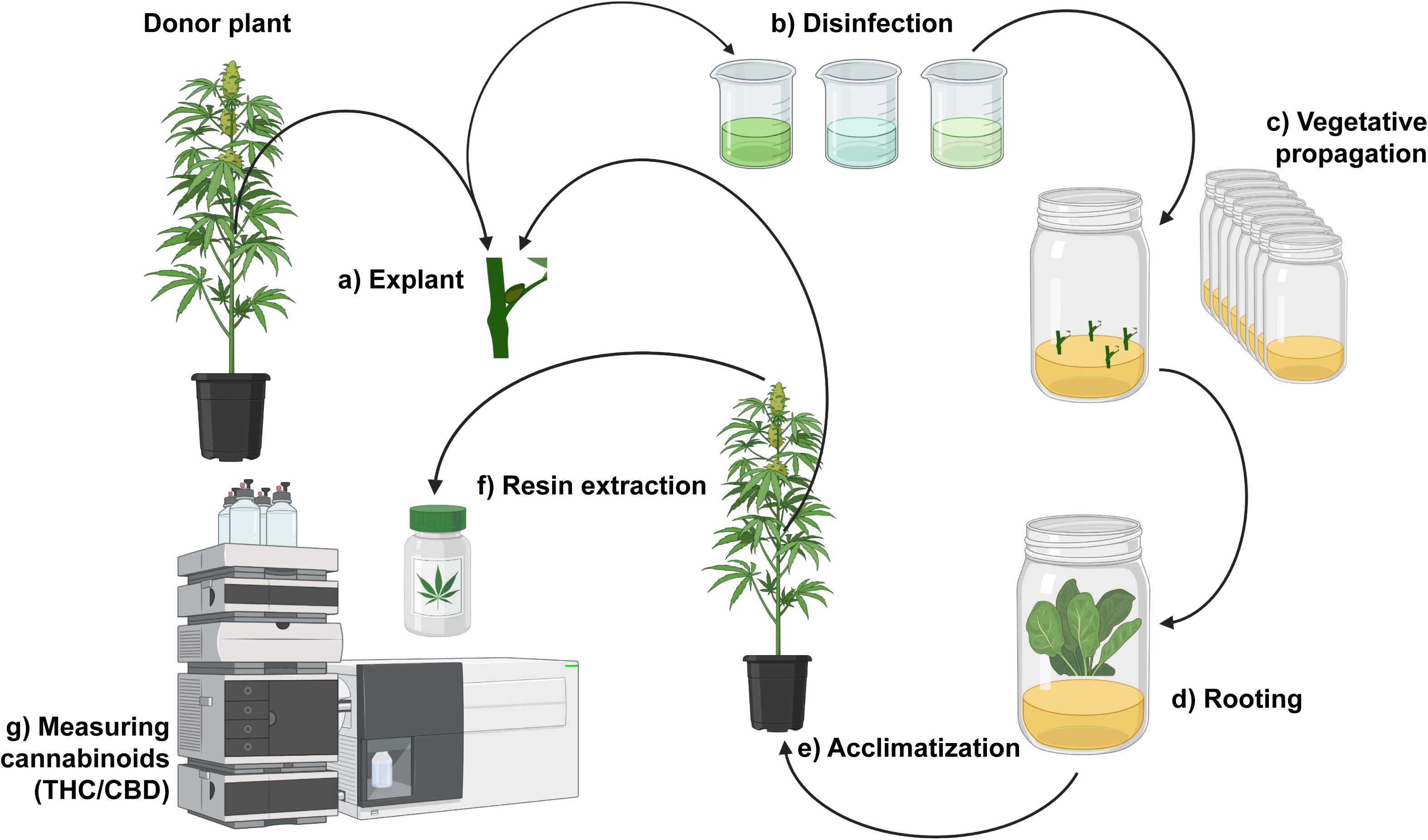
Schematic representation of the micropropagation protocol used to obtain cloned seedlings from a donor plant of the Charlie’s Dream variety of *Cannabis sativa*.

### 2.3. Disinfection

Explants were surface-disinfected by immersion in 80% ethanol for 1 minute with agitation, followed by treatment with 0.5% w/v sodium hypochlorite solution (Anedra) with 0.1% Tween 20 (Biopack) for 20 minutes. Subsequently, they were rinsed three times (5 minutes each) with sterile distilled water (Fig. 1b), dried on sterile filter paper in a laminar flow hood, and transferred to the corresponding culture media.

### 2.4. Culture media

MS medium (PhytoTechnology Laboratories) contained the composition of macro and micronutrients described by [11], sucrose (30 g/L, Biopack), and agar-agar (8 g/L, Britania). MSS was supplemented with nicotinic acid (0.01 g/L, Sigma), pyridoxine (0.01 g/L, Sigma), and thiamine (0.01 g/L, Sigma). Both media were adjusted to pH 5.8 and autoclaved for 20 minutes at 121°C. The plant growth regulators evaluated for vegetative growth were thidiazuron (TDZ, 0.5, 1, and 2.5 µM, Sigma-Aldrich) and benzylaminopurine (BAP, 0.5, 1, and 2.5 µM, Sigma-Aldrich) (Fig. 1c). Culture media were dispensed into sterilized glass jars, each initially containing four to five explants. Following the first subculturing, two explants were maintained per jar.

### 2.5. Growth of axillary and apical buds

The explants were incubated in culture chambers with a 16-h photoperiod, LED lights with a photon flux density of 200 µmol m^-2^ s^-1^, and a temperature range of 23°C±2°C, and the explants were repotted every 15 days. At 30 days post-initiation of micropropagation (dpim), gibberellin (7 µM, Sigma-Aldrich) was added to the MS medium containing the respective growth regulators, or the explants were exposed to red LED light (Interelec, 56 µmol m^-2^ s^-1^, Figure S1) to promote shoot elongation. Shoot length, number of shoots per explant, and foliar area were quantified from digital images using the ImageJ software. These parameters were assessed at 15, 30, and 45 dpim (Fig. 1c).

### 2.6. Rooting promotion and acclimatization

For *in vitro* rooting promotion, shoots were transferred to MS and MSS medium containing naphthaleneacetic acid (NAA, 2.5 and 5 µM, Sigma-Aldrich) or indolbutyric acid (IBA, 2.5 and 5 µM, Sigma-Aldrich) as rooting inducers (Fig. 1d). Additional media formulations were tested, containing ½ and ¼ strenght micro and macronutrients as described by [11], along with sucrose (30 g/L) and agar-agar (8 g/L), in the presence and absence of the auxins NAA (2.5 and 5 µM) or IBA (2.5 and 5 µM). For *ex vitro* rooting, shoots were transferred to pots filled with 1:1 perlite:soil substrate.

Prior to transplantation, the basal ends of the shoots were immersed in a solution NAA or IBA solution (2.5 or 5 µM) for 30 seconds (Fig. 1f). Root length was measured after four weeks for each treatment.

### 2.7. Resin extraction and cannabinoid analysis

For resin extraction, 5 g of dried inflorescences stored at -80°C were used. Without prior grinding, 250 mL of 96°C ethanol (also stored at -80°C) was added to each sample and stirred for 15 minutes. The extract was filtered twice using filter paper. The solvent removal was initially performed in a water bath at 45 °C and the residual solvent was evaporated in a SpeedVac concentrator at 45 °C. For cannabinoid analysis, resins were diluted in a 1:1 mixture of chloroform and methanol (CHClL:CHLOH). One microliter of each sample was injected into an Agilent 7890B gas chromatograph coupled to an Agilent 5977A mass spectrometer, equipped with an HP-5MSUI capillary column (30 m × 0.25 mm i.d., 0.25 μm film thickness). Mass spectra were identified by comparison with the NIST 2011b library. Quantification of CBD and Δ9-THC was carried out using calibration curves generated with certified standards from Cayman Chemical.

### 2.8. Statistical analysis

Statistical data analysis was performed using a two-way ANOVA followed by Tukey’s multiple comparison test. Differences were considered statistically significant at p < 0.05. GraphPad Prism version 8.0.2 was used for the analysis.

## 3. Results

### 3.1. Effect of supplementation with vitamins nicotinic acid, pyridoxine, and thiamine

The effect of the vitamins nicotinic acid, pyridoxine, and thiamine (MSS) on shoot length, number of shoots per explant, and foliar area was assessed in the presence of the growth-promoting hormones TDZ and BAP at three different concentrations (0.5, 1, and 2.5 µM).

The results indicated that increasing TDZ concentrations in both MS and MSS media significantly reduced the number of shoots per explant and the foliar area. The supplementation with any of the three vitamins did not enhance any of the parameters evaluated (Fig. 2A). At 15 dpim, explants cultured in MSS medium supplemented with 0.5 µM TDZ showed fewer shoots per explant than those grown in MS with the same concentration of TDZ (Fig. 2A). Higher concentrations of TDZ also negatively affected the foliar area of the explants (Fig. 2A). At 30 dpim, explants in MSS with 2.5 µM TDZ showed a smaller foliar area compared to those in MS mediunm at any TDZ concentration. It is worth mentioning that the use of 2.5 µM TDZ in both media-induced callus formation (data not shown).

**Figure 2.**
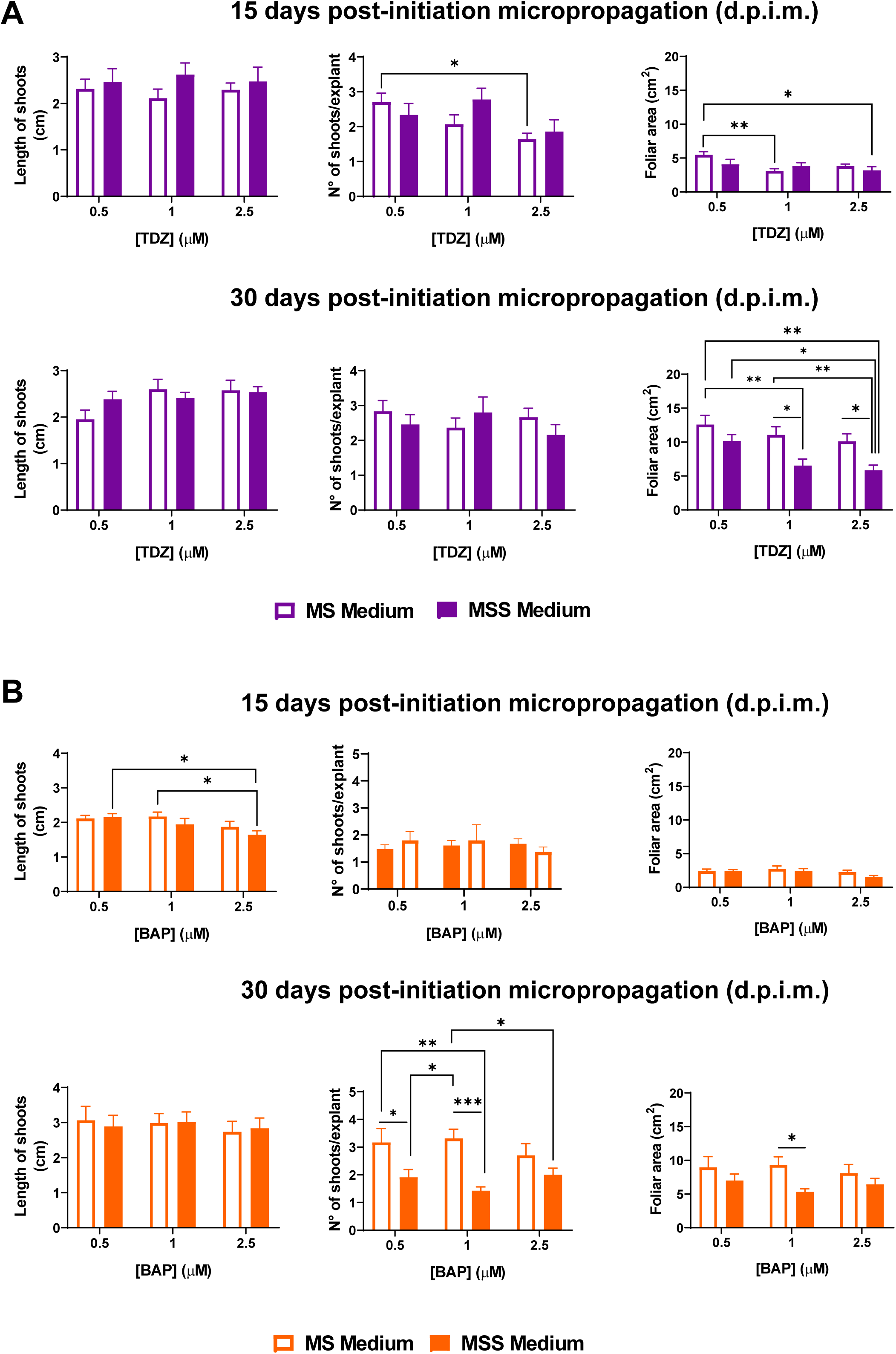
Analysis of the effect of the nicotinic acid, pyridoxine, and thiamine vitamins on the shoot length, number of shoots per explant, and foliar area. A) Effect of vitamin supplementation combined with the growth-promoting hormone TDZ at 15- and 30-day post-initiation micropropagation. B) Effect of vitamin supplementation combined with the growth-promoting hormone BAP at 15- and 30-day post-initiation micropropagation. Bars represent mean + SEM. Two-way ANOVA followed by Tukey’s multiple comparison test was used. * p < 0.05; ** p < 0.01; *** p < 0.001.

In contrast, the addition of BAP to MS and MSS media enabled the evaluation of vitamin supplementation effects on shoot number and foliar area (Fig. 2B). At 30 dpim, explants cultured in MS with 1 µM BAP exhibited a higher number of shoots per explant than those in MSS medium with BAP at any concentration (Fig. 2B). Likewise, foliar area was greater in explants from MS with 1 µM BAP compared to those from MSS with the same BAP concentration. Overall, vitamin supplementation negatively affected explant development when BAP was used as a vegetative growth-promoting hormone.

### 3.2. Effect of TDZ and BAP hormones

Since the MSS medium did not enhance any of the three parameters evaluated, a comparison between the growth-promoting hormones TDZ and BAP was performed using the MS medium. At 15 dpim, explants cultured in MS with 0.5 µM TDZ showed a more shoots compared to explants grown in MS with 0.5 µM BAP medium (Fig. 3A). Similarly, explants treated with 0.5 µM or 2.5 µM TDZ exhibited greater foliar area development compared to those treated with the same concentrations of BAP (Fig. 3A). However, at 30 dpim, the explants grown in the presence of BAP exhibited improvements across all three parameters evaluated, reaching values comparable to those obtained with TDZ (Fig. 3A, Fig. S2).

**Figure 3.**
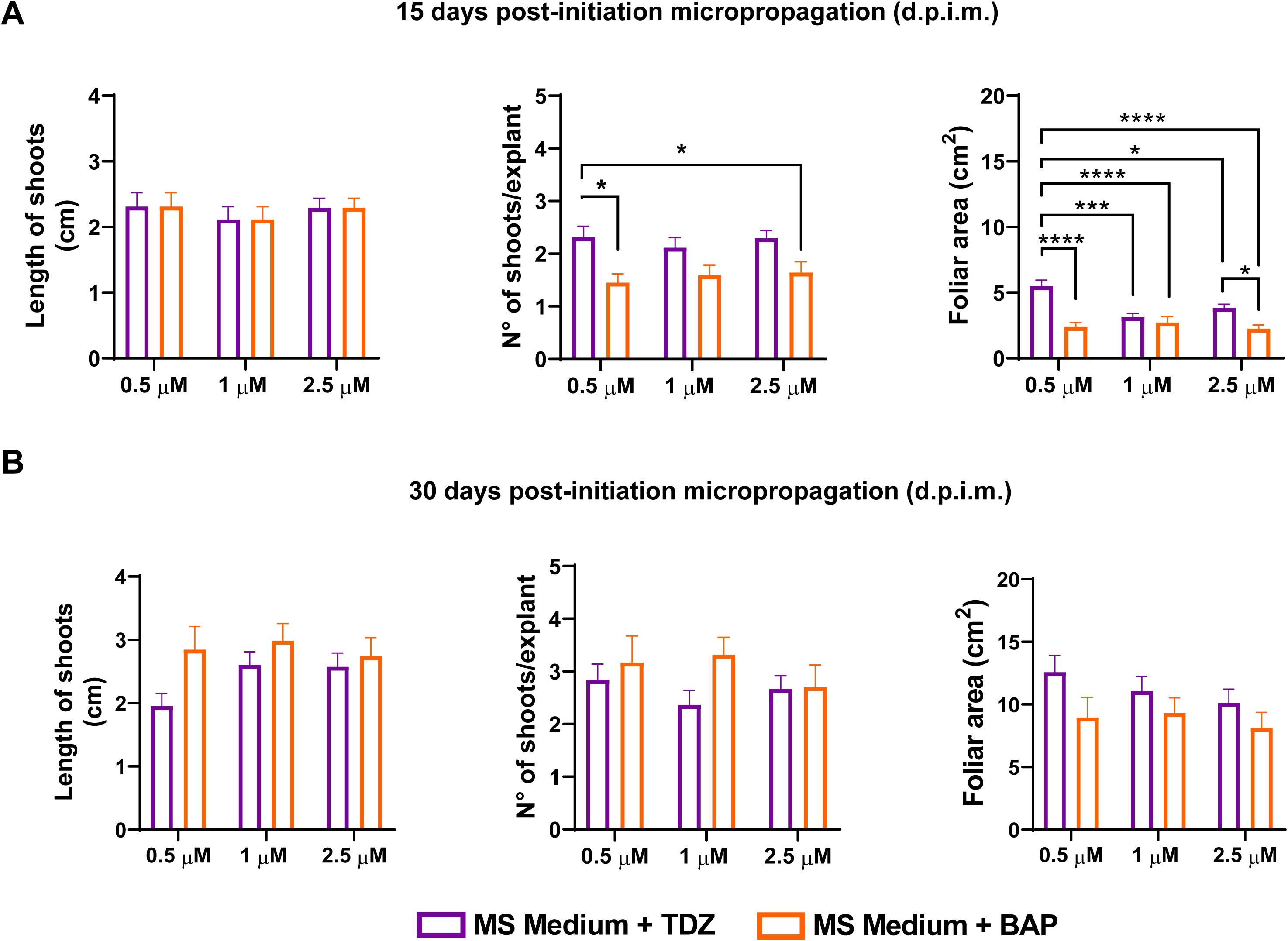
Analysis of the effect of the growth-promoting hormones TDZ and BAP on shoot length, number of shoots per explant, and foliar area at 15 (A) and 30 days (B) post-initiation micropropagation. Bars represent mean + SEM. Two-way ANOVA followed by Tukey’s multiple comparison test was used. * p < 0.05; *** p < 0.001; **** p < 0.0001.

### 3.3. Effect of Gibberellin and LED red light

Given that no significant increase in shoot length was observed between 15 and 30 dpim, MS + growth-promoting hormones medium was supplemented with gibberellin, and its effect was compared with that of incubation under red light, as both treatments have been reported to influence shoot elongation under *in vitro* conditions [12,13]. The results showed that both gibberellin supplementation and red-light exposure enhanced shoot length, but only when combined with BAP at 0.5 or 1 µM (Fig. 4A). In contrast, when these treatments were combined with TDZ (0.5 or 1 µM), shoot length was negatively affected (Fig. 4A, Fig. S3).

**Figure 4.**
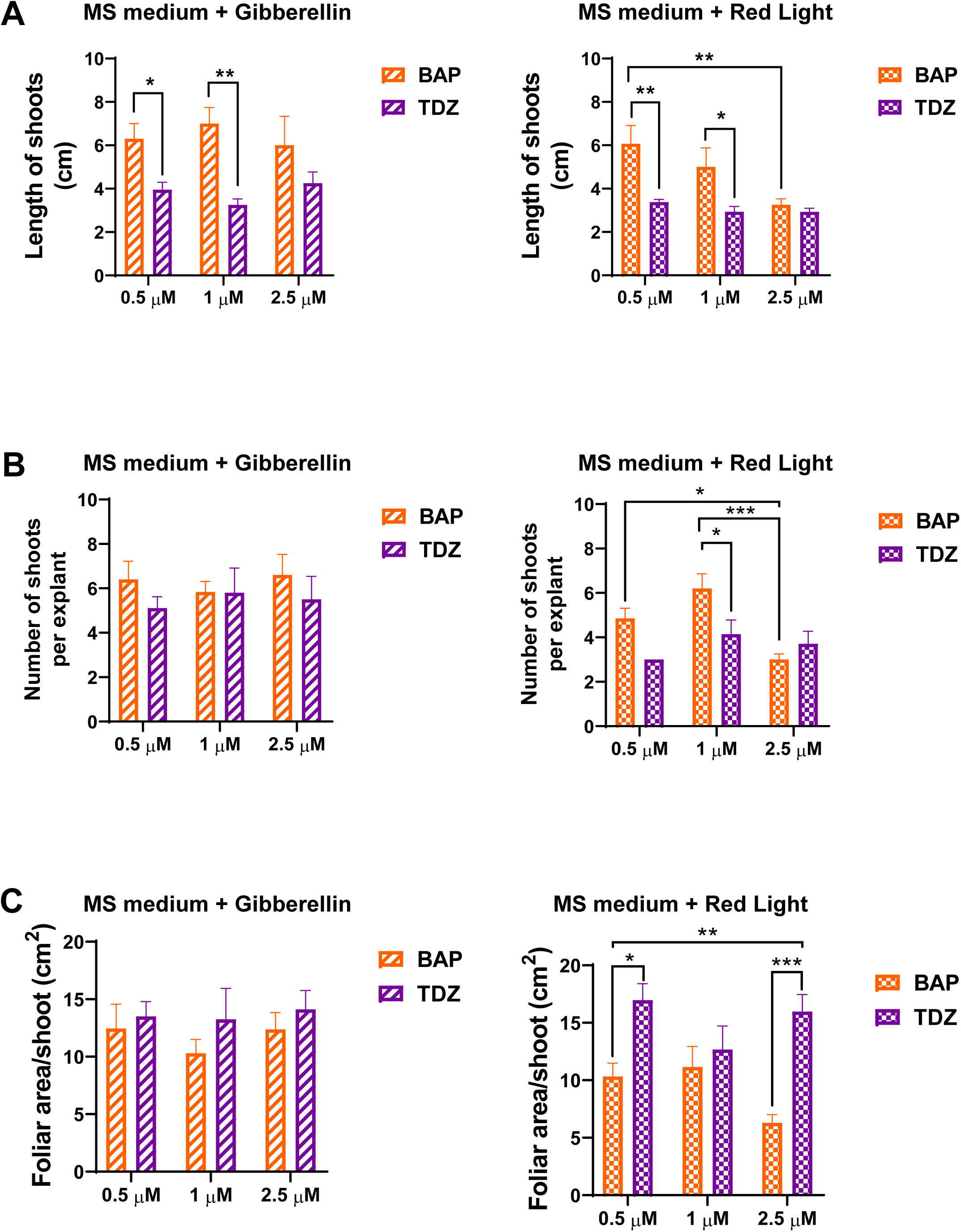
Analysis of the effect of the gibberellin hormone and red LED light on shoot length (A), number of shoots per explant (B), and foliar area per shoot (C) at 45 days post-initiation of micropropagation. Bars represent mean + SEM. Two-way ANOVA followed by Tukey’s multiple comparison test was used. * p < 0.05; *** p < 0.001; **** p < 0.0001.

An increase in the number of shoots per explant was observed with both gibberellin and red-light treatments (Fig. 4B), compared to the previously described results (Fig. 3B). However, explants cultured in the presence of red light and TDZ produced fewer shoots than those treated with red light and BAP (Fig. 4B). Moreover, combining 2.5 µM BAP with red LED light negatively affected both shoot length and the number of shoots per explant (Fig. 4A, B). Finally, treatments with red light and gibberellin promoted an increase in the foliar area (Fig. 4C). However, no significant differences were observed between the two hormones when shoots were treated with gibberellin (Fig. 4C). In contrast, treatment with red light and TDZ significantly increased foliar area compared to red light and BAP (Fig. 4C).

### 3.4. Effect of the hormones IBA and NAA

The rooting of the obtained shoots was evaluated through two treatment groups. Rooting was assessed under *in vitro* culture conditions or through *ex vitro* rooting. For *in vitro* root induction, shoots were randomly placed in different glass flasks containing MS, MS½, and MS¼ medium, which were either supplemented or not with the root growth-promoting hormones NAA (2.5 µM or 5 µM) or IBA (2.5 µM or 5 µM); however, rooting was not induced under any of the tested conditions. For *ex vitro* rooting, 30-dpim shoots were transplanted into pots after being pre-treated or not with NAA or IBA at the previously mentioned concentrations (Fig. 5). Greater root length was observed in shoots that were pre-treated with 2.5 µM IBA compared to control shoots (without hormone treatment) and those pre-treated with NAA at either concentration (Fig. 5).

**Figure 5.**
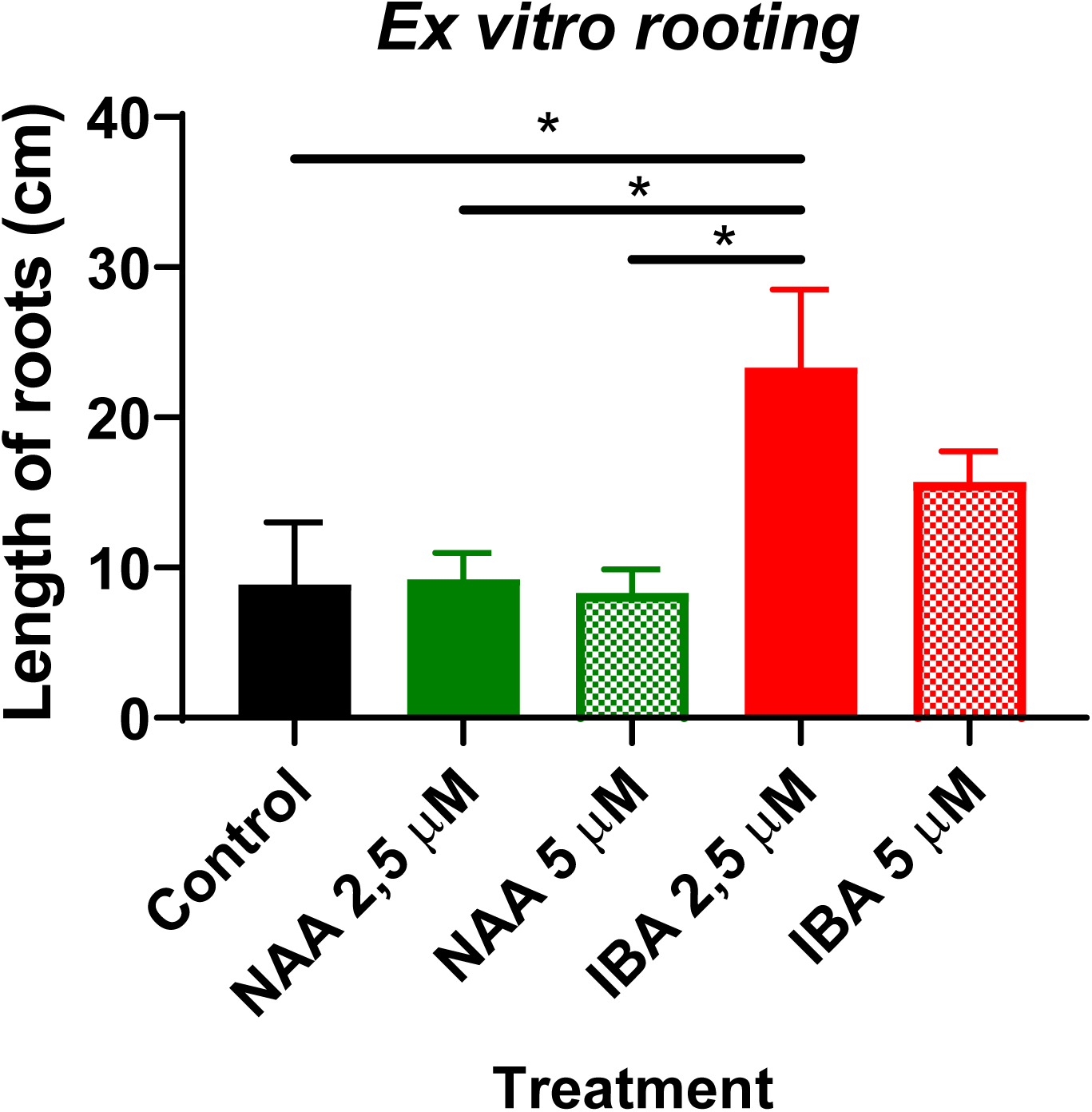
Effect of IBA and NAA on root length. Bars represent mean + SEM. Statistical analysis was performed using two-way ANOVA followed by Tukey’s multiple comparison test. * p < 0.05; *** p < 0.001; **** p < 0.0001.

### 3.5. Analysis of cannabinoid profile

Resins extracted from the flowers of two donor plants, four micropropagated plants (G1) derived from one of the donor plants (donor plant 2), and eighteen micropropagated plants obtained from the four G1 plants (G2) were used to quantify CBD and Δ^9^-THC levels (Table 1). In addition, the relative percentages of CBD, Δ^9^-THC, and other cannabinoids were determined in the resin from donor plant 2 and from the micropropagated plants (G1 and G2) (Table 2). The results showed that the CBD/Δ^9^-THC ratio and the relative percentages of cannabinoids remained within similar values between the donor plant and the micropropagated plants derived from it. Likewise, markedly different CBD/Δ^9^-THC ratios were observed between the samples from both donor plants, suggesting that greater variability in cannabinoid content between plants of the same chemotype would be generated by seed propagation. It is worth noting that a resin yield of 15 ± 2 g per 100 g of dried flowers (n = 18) was obtained under our flowering conditions and through a single resin extraction procedure.

## 4. Discussion

In this work, the effect of vitamin supplementation (nicotinic acid, pyridoxine, and thiamine) in MS medium with TDZ and BAP on the growth and development of axillary and apical buds from the Charlie’s Dream variety of *C. sativa* (chemotype III, THC<<CBD) was evaluated. The number of shoots per explant, shoot length, and the foliar area were analyzed as development parameters of vegetative development. Differentiated effects were observed in both the short and long term, providing a more detailed understanding of how these hormones and vitamins influence cell proliferation and plant vegetative development.

At higher concentrations of TDZ, a negative effect on growth was observed, with a significant reduction in the number of shoots per explant. These effects may be explained by the capacity of TDZ to induce cell dedifferentiation at high concentrations, thereby inhibiting shoot proliferation and promoting callus formation [14–17]. The negative effect of TDZ was not counteracted by the use of a vitamin-supplemented medium (MSS), as vitamin supplementation did not enhance cell proliferation or differentiation. On the other hand, BAP treatment also resulted in variable responses depending on the concentration used. Our results agree with previous studies, where a concentration of 1 µM BAP was reported to be the most suitable for promoting shoot proliferation in *C. sativa* [12,14,18,19]. However, explants grown in the MSS medium exhibited reduced foliar development and shoot number, suggesting that the vitamins in MSS may have interfered with the action of BAP on cell proliferation. This inhibitory effect could be related to a metabolic interaction between vitamins and hormones in the explants, creating an unfavorable environment for explant development. These findings are particularly relevant for the optimization of micropropagation protocols, as they indicate that vitamin supplementation may not be necessary when appropriate concentrations of BAP are used. The behavior of BAP may be associated to its role in promoting cell differentiation and organogenesis, facilitating balanced growth throughout developmental phases [18]. These findings also highlight the importance of adjusting growth-promoting hormones (in this case, BAP and TDZ) concentrations to prevent adverse effects and to maximize cell proliferation without inducing dedifferentiation.

Gibberellin has been used to stimulate shoot elongation and organ differentiation in micropropagation protocols [20,21]. In addition to hormonal regulation, environmental factors such as light quality are also known to affect plant development [22,23]. In the present study, shoot elongation was enhanced by gibberellin supplementation and red-light exposure, particularly when explants were cultivated in MS medium supplemented with BAP at 0.5 or 1 µM. These findings agree with previous reports indicating that red light and gibberellin exposure promote shoot elongation under *in vitro* conditions [24,25]. However, when these treatments were combined with TDZ, shoot elongation was negatively affected, consistent with the inhibitory action of TDZ on shoot formation and differentiation [2,15]. Furthermore, explants exposed to TDZ under red light showed a reduced number of shoots per explant compared to those exposed to red light with BAP, reinforcing the idea that TDZ may hinder shoot proliferation. The reduced shoot elongation observed when using 2.5 µM BAP combined with red light further emphasizes the need for precise hormone concentration optimization to support *in vitro* plant growth and development.

Concerning rooting, our results point to this stage as one of the main bottlenecks in *C. sativa* micropropagation. The effects of the hormones IBA and NAA on the rooting of shoots derived from axillary and apical buds of the Charlie’s Dream variety were evaluated under both *in vitro* and *ex vitro* rooting conditions. Under *in vitro* conditions, root development was not achieved under any treatment tested, and the shoots remained at the callus stage. Although difficulties in achieving *in vitro* rooting of *Cannabis sativa* in full-strength MS medium have been reported [16,26], rooting has been observed in ½ MS medium. In our case, however, shoot necrosis occurred within one week of culture in both half- and quarter-strength MS medium. Moreover, although red LED light irradiation has been shown to improve *in vitro* rooting in hard-to-propagate species [27], no root development was under this treatment either. In contrast, under *ex vitro* conditions, greater root length was observed when shoots were pretreated with 2.5 µM IBA compared to untreated controls or those treated with NAA. This outcome is consistent with findings from other studies, who have found that IBA is more effective than NAA in inducing rooting in different cannabis varieties [2,14,28,29]. The success of ex vitro rooting appears to have been favored by the direct contact of shoots with the substrate.

Finally, the cannabinoid profile of micropropagated plants was compared with that of the donor plants. The results indicated that the CBD/Δ^9^-THC ratio and the relative percentages of other therapeutically relevant cannabinoids remained stable across two generations of micropropagation. Although only the short-term cannabinoid profile was assessed in this study, future work will be necessary to determine whether the cannabinoid profile remains stable over successive micropropagation cycles.

## 5. Conclusion

The present study demonstrated that the growth and development of axillary and apical buds from the *C. sativa* Charlie’s Dream variety are significantly influenced by the concentrations of TDZ and BAP, as well as by the vitamin content in the culture medium. The findings underscore the importance of optimizing hormonal and nutritional conditions to promote effective shoot proliferation and elongation during *in vitro* propagation. Moreover, clonal micropropagation of Charlie’s Dream variety was validated as a reliable strategy for maintaining the chemical stability of cannabinoid profiles across generations, a critical factor for ensuring product consistency in the medicinal cannabis industry. Given the challenges observed during the rooting stage, particularly under in vitro conditions, future investigations should consider the role of soil microbiota, particularly plant growth-promoting bacteria (PGPB), in contributing to *ex vitro* rooting through phytohormone production or other growth-stimulating mechanisms [30].

## Funding

This study was supported by ex Ministerio de Ciencia, Tecnología e Innovación (proyecto 14A).

## Author Contributions

M.G. and G.Y.: Experimental design, data analysis, and figure design; A.S.: Experimental design and data analysis. M. H., E. B., M. D. L., and L. S.: Chemical analysis and data analysis. E.P.: Experimental design; M. C.: Experimental design, data analysis, manuscript writing, and editing. All authors read the manuscript and contributed to the writing and final editing.

## Institutional Review Board Statement

Neither humans nor animals were utilized in this study.

## Informed Consent Statement

Not applicable

## Credit authorship contribution statement

Micaela Gonzalez and Gabriel Yañuk: Writing – original draft, Visualization, Methodology, Investigation, Formal analysis, Conceptualization. Analía Sannazzaro: Writing – review & editing, Supervision, Formal analysis, Conceptualization. Eliane Perez: Methodology, Visualization. Estefanía Butassi, Melina Di Liberto, Laura Svetaz: Ignacio Cabezudo and Mónica Hourcade; Writing-review & editing, Methodology, Formal analysis, Supervision. Marina Clemente: Writing – original draft, Investigation, Formal Analysis, Conceptualization, Supervision.

## Declaration of Competing Interest

None of the authors has any conflicts of interest.

## Acknowledgments

The authors thank Patricia Uchiya (CIC-pba) and Carlos Alberici (CONICET) for their technical assistance and the GC-EM service of the Facultad de Ciencias Bioquímicas y Farmacéuticas (Universidad Nacional de Rosario, Argentina).

## Data availability

Data will be made available on request.

## Legends of the supplementary figures

**Figure S1.** Spectral irradiance of the red LED light array, measured under the same experimental conditions used during the shoot elongation phase, using a USB2000 spectroradiometer (Ocean Optics) equipped with a fiber optic cable and a Teflon diffuser.

**Figure S2.** Representative photographs of CharlieLs Dream explants grown in the presence of BAP and TDZ (0.5, 1, and 2.5 µM) after 30 days of culture.

**Figure S3.** Influence of gibberellin and red LED light on shoot elongation. Representative photographs of CharlieLs Dream explants grown for 15 days in the presence of BAP or TDZ (0.5, 1, and 2.5 µM) in combination with gibberellin (7 µM) or red light.

## Notes

### Competing Interest Statement

The authors have declared no competing interest.

