## Supplementary figures and images for "Optimization of a *Cannabis sativa* micropropagation protocol (chemotype III) to preserve its cannabinoid profile"

### Figure S1

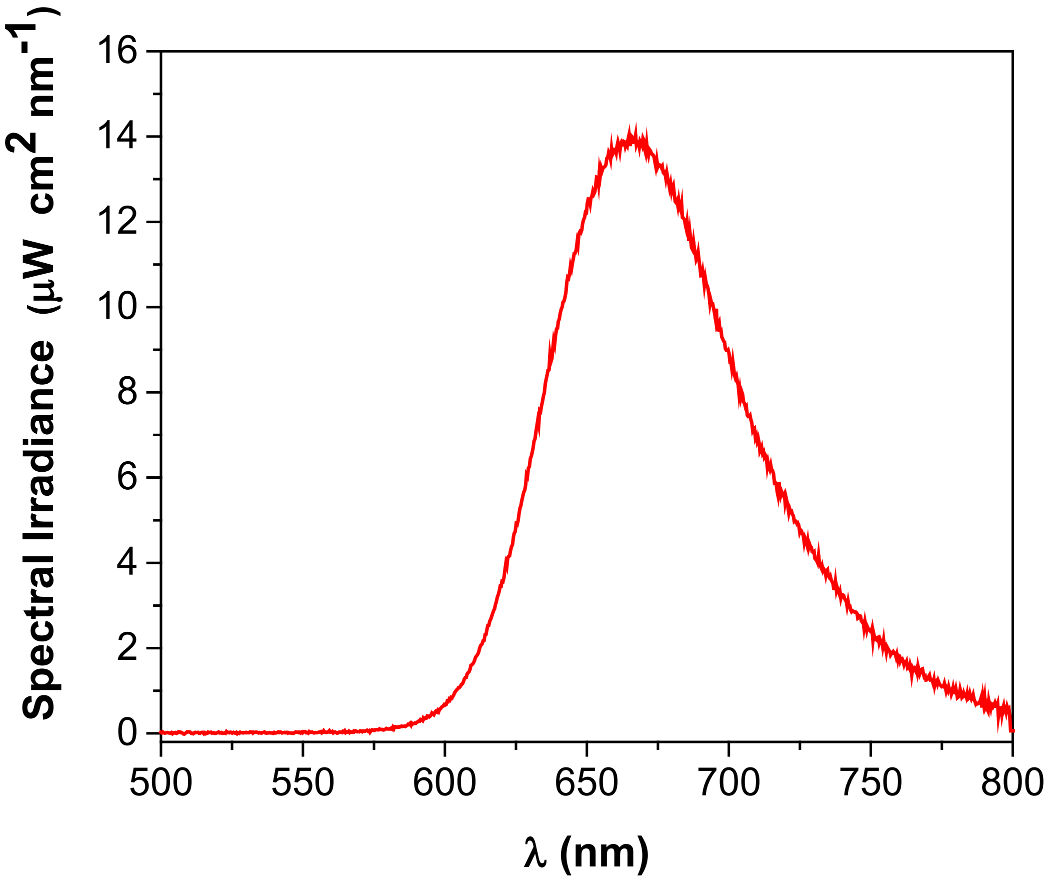

### Figure S2

Fig. S2

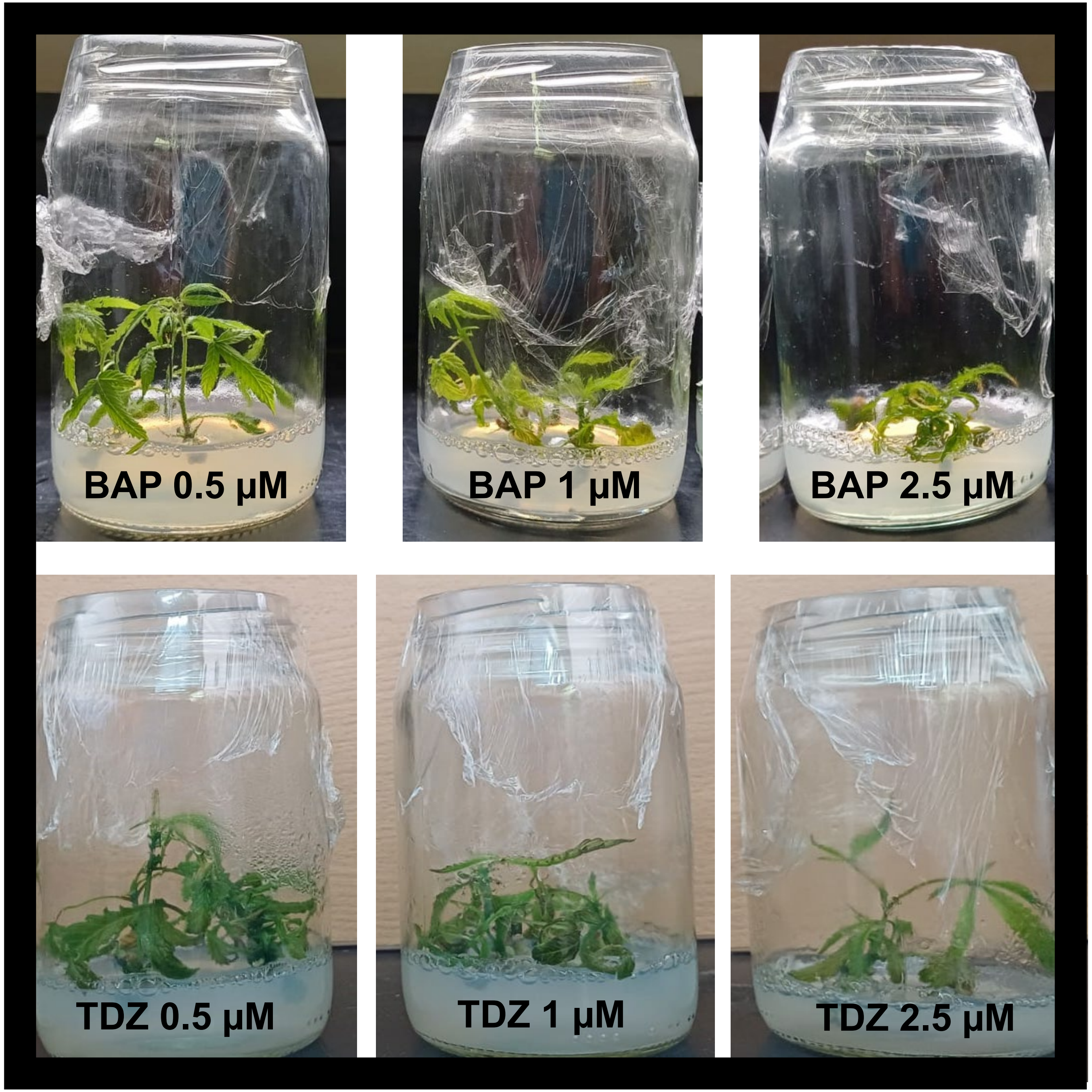

### Figure S3

Fig. S3

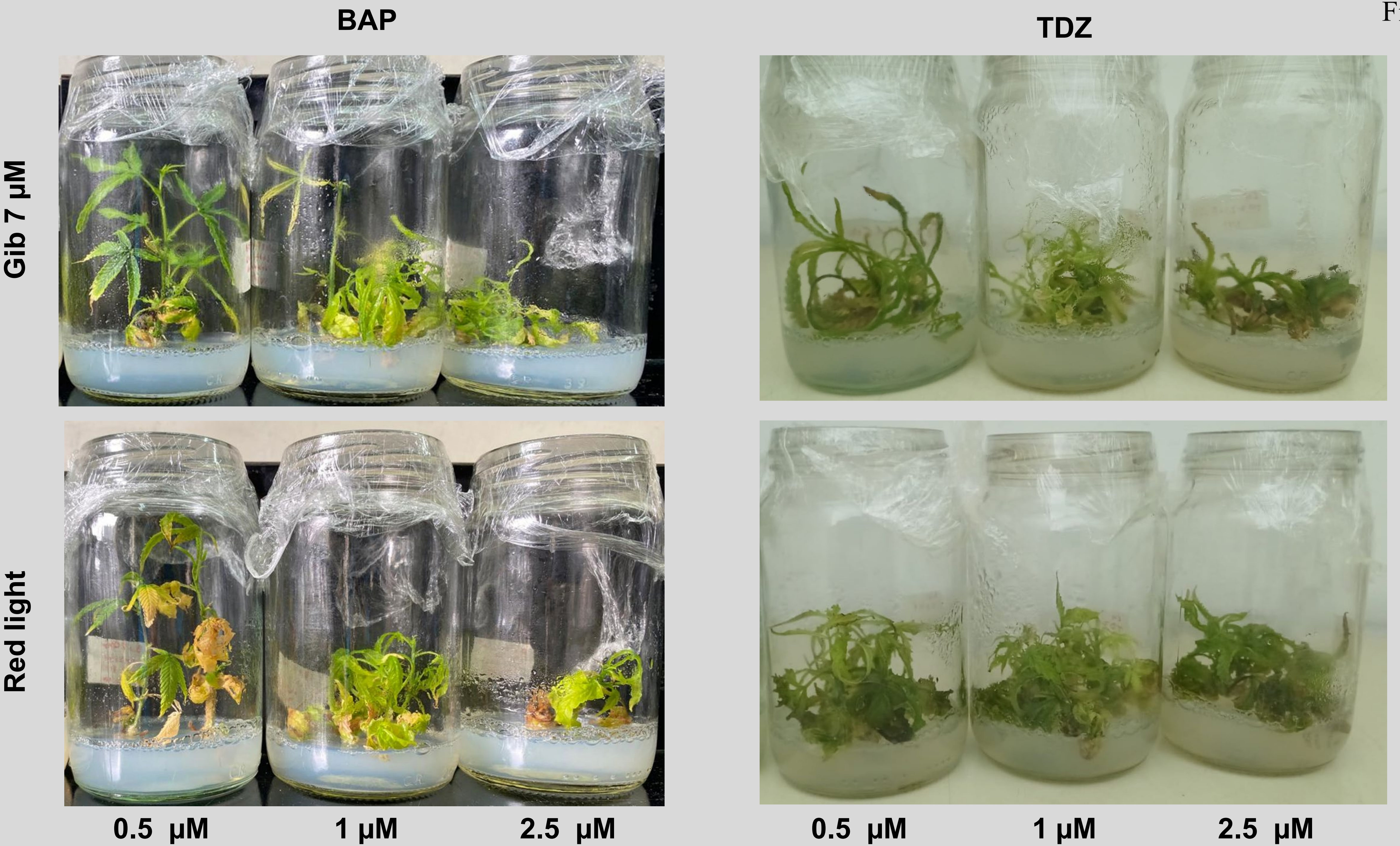
